# Structured Multimodal Deep Learning improves Genomic Prediction in Future Environments

**DOI:** 10.1101/2025.09.05.674546

**Authors:** Aike Potze, Fred van Eeuwijk, Ioannis N. Athanasiadis

## Abstract

The development of prediction models for phenotypes as functions of genetics and environmental inputs is a long-standing challenge in genetics and plant breeding. Deep neural networks form a promising approach to this task, due to their capacity to approximate nonlinear biological processes. Despite initial expectations, recent studies have found deep neural networks under-performing in comparison to linear methods, even for continent-scale trial datasets. We attribute this to several failure modes of deep learning, including *greedy learning*, the tendency of deep neural networks to over-emphasize a single type of input data. As a solution, we present the Structured Interaction Neural Network (SINN), which combines statistical decomposition of genetic, environmental and interaction effects with deep neural networks. SINN dissects phenotype prediction into isolated component modeling tasks, revealing poor generalization of learned representations to new environments to be the main limitation for both prediction of genotype-by-environment interactions and yield prediction overall. By balancing model complexity and regularization per component, we reach competitive performance on yield prediction in the next cycle of a maize multi-environment trial, including both new genotypes and new environmental conditions. SINN achieved a higher accuracy (0.63) than BLUP-based methods (0.43) and a neural network from previous literature (0.48), and surpassed the top-performing models in a public benchmark dataset with a lower RMSE (2.41 Mg/ha versus 2.46 Mg/ha, with mean yield of 9.51 Mg/ha) and higher genetic correlation (0.38 versus 0.36). By combining statistical genetics and modern deep learning, SINN enables accurate, modular and scalable genomic prediction in new environments.

## Introduction

Climate change is shifting local environmental conditions, which increasingly threatens global crop yields (Kang *et al*. 2009). A major focus of modern plant breeding is the mitigation of these effects of climate change on crop performance (Chapman *et al*. 2012). Climate change can also directly affect the effectiveness of breeding programs, as it can reduce alignment between a multi-environment trial (MET) and the target population of environments (TPE) (Cooper *et al*. 2023). To address this challenge, new methods are needed that predict how new crop genotypes and environments lead to a desired phenotype. As this task forms an extension of genomic prediction (GP) (Meuwissen *et al*. 2001) to new environments, we refer to this prediction task as genomic-enviromic prediction (GEP).

Interest in GEP has surged in recent years, inspiring a diverse set of novel methods. Proposed statistical methods include factorial regression on environmental covariates (ECs) (Heslot *et al*. 2014), linear mixed models using enviromic relationship kernels (Jarquín *et al*. 2014), linear and linear mixed models using grouping of environments in envirotypes (Xu 2016) and mega-scale linear mixed models (Hu *et al*. 2025). Additionally, various methods incorporating process-based crop growth models (Jullien *et al*. 2011; Technow *et al*. 2015; Cooper *et al*. 2016; Rincent *et al*. 2019) and machine learning models (Westhues *et al*. 2021; Fernandes *et al*. 2024; He *et al*. 2025) have been proposed. Recently, deep learning (DL) has emerged as a promising modeling paradigm due to its capacity to model nonlinear relationships (Goodfellow *et al*. 2016), predictable improvements in predictive performance with dataset size (Hestness *et al*. 2017), and flexibility in combining diverse data (Ngiam *et al*. 2011).

Multimodal deep learning (MMDL), a category of DL methods combining multiple types (modalities) of data, can integrate enviromics, phenomics and other omics into genomic prediction (Montesinos-López *et al*. 2024). Adoption of MMDL for GEP has been relatively limited (Jubair and Domaratzki 2023), with mixed success in case studies for wheat (Montesinos-López *et al*. 2019; Guo *et al*. 2020; Sandhu *et al*. 2021; Jubair *et al*. 2023), corn (Montesinos-López *et al*. 2018; Khaki and Wang 2019; Washburn *et al*. 2021; Kick *et al*. 2023) and barley (Måløy *et al*. 2021). Despite the widespread optimism and the isolated improvements reported in aforementioned works, open benchmarking of models for GEP on a continent-scale MET found MMDL methods not to be competitive with machine learning and statistical methods (Washburn *et al*. 2025). Considering the massive scale of this data, with over 100,000 measured plots (Lima *et al*. 2023), this suggests that current MMDL approaches are ineffective at GEP for even the largest of current METs.

Current MMDL research tempers the aforementioned optimism: multimodal models do not necessarily improve over unimodal approaches, and are not guaranteed to learn any interactions. Multimodal architectures are more complex than unimodal architectures, which harms performance through *over-fitting* (Wang *et al*. 2020). The rate of this overfitting is modality-specific, which can contribute to *greedy learning*, the tendency to over-rely on a single dominant modality while underfitting on additional modalities (Cadene *et al*. 2019; Huang *et al*. 2022; Peng *et al*. 2022; Wu *et al*. 2022; Du *et al*. 2023; Makino *et al*. 2023; Zhang *et al*. 2024). Counterintuively, while multimodal models are often motivated from the prediction of interactions, they can function like the sum of unimodal models (Hessel and Lee 2020; Wörtwein *et al*. 2022), which indicates *underutilization of interactions*. Clearly, multimodal learning introduces problem-specific failure mechanisms. Unfortunately, it remains an open question which specific failure mechanisms apply to GEP.

To avoid these issues we introduce the Structured Interaction Neural Network (SINN), a new framework which enables the fitting of MMDL models for GEP. It contains four steps: first we utilize MET structure and statistical models to decompose phenotype into unimodal components (genetic, environmental) and multimodal components (genotype-environment interactions). These separated signals are then used to optimize unimodal DNNs for genomic prediction of the genetic component and enviromic prediction of the environmental component. Next, the unimodal DNNs are used to initialize a multimodal DNN that predicts the genotype-environment interactions. Finally, the predictions of all three DNNs are summed to obtain predictions for new phenotypes. By separating unimodal and multimodal signals, this approach sidesteps greedy learning and underutilization of interactions. Furthermore, the structured optimization strategy enables us to finely tune the complexity of each component DNN to each modality, reducing overfitting.

We demonstrate the effectiveness of SINN by predicting the yield of new maize hybrids in the next year of a large-scale MET in the United States of America (Lima *et al*. 2023), collected and shared through by the Genomes-to-Fields (G2F) Initiative (Lawrence-Dill *et al*. 2019). As we directly follow the benchmarking protocols of a 2022 competition with identical data (Washburn *et al*. 2025), we are able to fairly compare SINN against the state-of-the-art across 124 models submitted to the competition.

In short, we present SINN, a decomposition-based neural network framework for genomic-enviromic prediction of phenotype that:

i. Improves prediction of yield for new maize genotypes in new environments
ii. Increases utilization of genetic information of DNNs
iii. Simplifies the development of multimodal DNNs
iv. Supports targeted improvement of genotype-environment interaction prediction

## Materials and methods

In this section, we describe our approach to yield prediction for new genotypes and new environments. Figure 1 shows an overview of the approach. Let *i* = 1, 2, …, *n*_*g*_ denote the *n*_*g*_ crop genotypes and *j* = 1, 2, …, *n*_*e*_ denote the *n*_*e*_ growing environments. For each genotype *i* we define a vector of *d*_*g*_ genetic markers 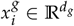, representing genetic variations across the crop genome. For each environment *j* we define a vector of *d*_*e*_ environmental covariates 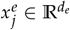, representing observations of weather, soil and management variables. Furthermore, we define the crop yield of genotype *i* in environment *j* as *y*_*ij*_ ∈R _≥0_. We then formalize the our objective as follows. Given a set of *n*_*s*_ training observations 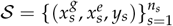, learn a mapping *f* from genetic markers 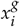 and environmental covariates 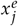 to crop yield *y* for any any genotype-environment combination (*i, j*).

**Figure 1.**
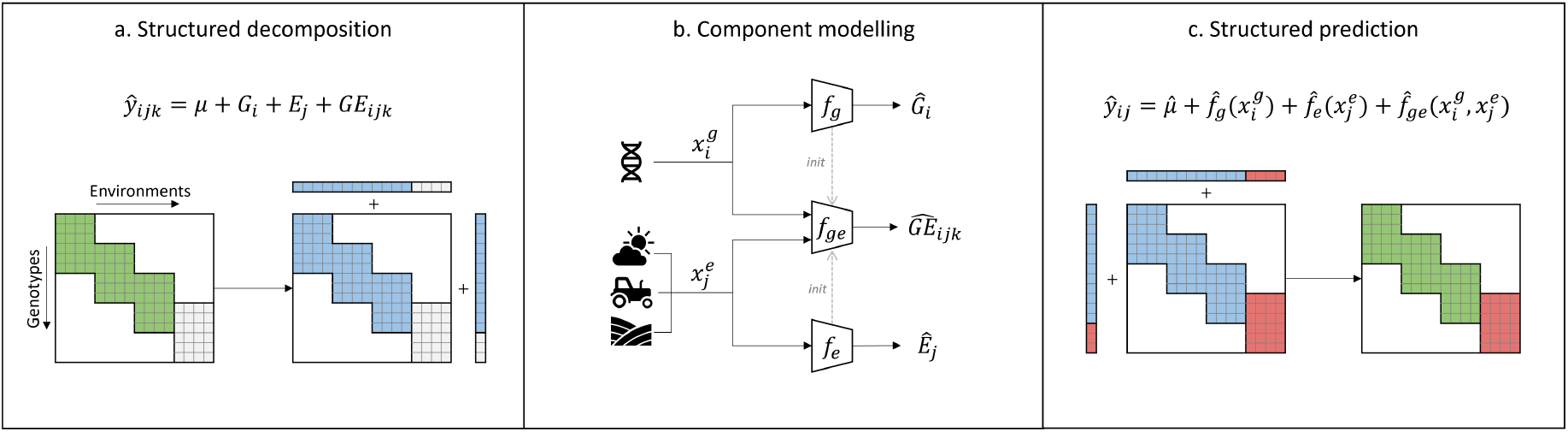
Overview of the proposed Structured Interaction Neural Network (SINN). a) Observed plot yields (green) are decomposed into yield components for genotypes, environments and interactions (blue). b) Estimated components are used to train three DNNs, which predict yield component from genetic markers 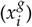 and environmental covariates 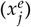, including weather, management and soil. The DNNs for genetic and environmental components are used to initialize the DNN for the interaction component, denoted by *init*. c) Trained DNNs predict each component for new genotypes and environments (red). Finally, DNN predictions are recomposed to predict the yield for unobserved plots in the next cycle of a multi-environment trial.

We define the quality of mapping *f* as the difference between predicted yield 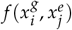 and observed yield *y* on a set of held-out samples. We distinguish between four prediction or cross-validation (CV) scenarios, depending on whether genotype *i* and environment *j* were previously observed in 𝒮:

1. **CV-X**: Observed genotypes, observed environments
2. **CV-G**: New genotypes, observed environments
3. **CV-E**: Observed genotypes, new environments
4. **CV-GE**: New genotypes, new environments

While all four scenarios serve useful applications, they are not equally conductive to the comparison of flexible predictive models. Repeated structures, such as previously observed genotypes, can be exploited as shortcuts by flexible models, including neural networks. This leads to an overestimation of the true predictive ability when comparing models on scenario CV-X, CV-G and CV-E. As such, we utilized the most stringent scenario CV-GE to compare the ability of various models to generalize yield prediction to new conditions.

### Introducing SINN

Direct approximation of yield can lead to overfitting (Wang *et al*. 2020), greedy learning (Wu *et al*. 2022) and underutilization of interactions (Hessel and Lee 2020). To avoid this issue, we do not directly approximate *y*, but instead define the following decomposed representation:

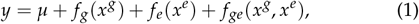

with components further defined as:

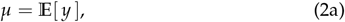

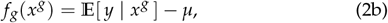

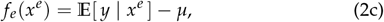

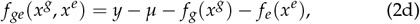

where *µ* represents the overall expectation of *y*, the two component functions *f*_*g*_ and *f*_*e*_, represent the unimodal contributions of the genetic and environmental features, and *f*_*ge*_ represents the contribution of genotype-by-environment interactions to *y*. Because genetic markers *x*^*g*^ and environmental covariates *x*^*e*^ are indexed by genotype *i* and environment *j*, respectively, we can approximate conditional expectation 𝔼 [*y* | *x*^*g*^] by 𝔼 [*y* | *i*] and conditional expectation 𝔼 [*y* | *x*^*e*^] by 𝔼 [*y* | *j*]. We subsequently estimate *µ*, 𝔼 [*y* | *i*] and 𝔼 [*y* | *j*] using the following linear fixed model:

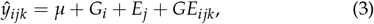

where *µ* represents the grand mean of *y, ŷ*_*ijk*_ represents the yield of genotype *i*, environment *j* and replicate *k, G*_*i*_ and *E*_*j*_ the estimated fixed effects of genotype and environment in *S*. Error term *GE*_*ijk*_ ∼ 𝒩 (0, *σ*^2^) then represents the residual genotype-by-environment interaction of the sample. We note that (3) represent a minimal decomposition. Nonetheless, we deliberately estimate *G*_*i*_ and *E*_*j*_ as fixed effects to avoid the adverse effects of shrinkage in downstream modeling (Holland and Piepho 2024).

Next, we utilize the fitted effects *Ĝ*_*i*_, *Ê*_*j*_ and residual *ĜE*_*ijk*_ from (3) as three sets of training labels to approximate mappings *f*_*g*_, *f*_*e*_ and *f*_*ge*_ from genetic features 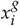 and environmental features 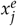:

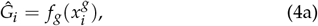

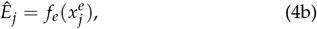

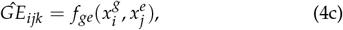

with *f*_*g*_, *f*_*e*_ and *f*_*ge*_ being three unique DNNs that are optimized separately. This effectively decomposes the prediction task into three component tasks: (4a) is a standard genomic prediction task, (4b) a standard enviromic prediction task, and (4c) a new interaction prediction task. The implementation details for each prediction task are described in *Model implementation* below. In order to integrate the learned representations from genomic prediction and enviromic prediction into interaction prediction, we initialize the parameters *θ*_*ge*_ of *f*_*ge*_ with the learned weights *θ*_*g*_ and *θ*_*e*_ of *f*_*g*_ and *f*_*e*_:

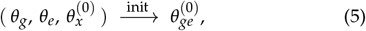

where the parameters of the interaction component of *f*_*ge*_, *θ*_*x*_ are initialized using any standard initialization. To predict *y* from genetic and environmental features for new genotypes and environments, one can finally substitute (4a), (4b) and (4c) in (3) to obtain

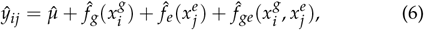

which directly estimates yield *y* for genotype *i* and environment *j* from genetic features 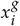 and environmental features 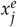. Together, _*i j*_ (2)-(6) form our proposed method, the Structured Interaction Neural Network (SINN). SINN utilizes the known structure of the training set to *𝒮* decompose *y*_*s*_ into independent label sets for genetic (*G*_*i*_), environmental (*E*_*j*_) and interaction effects (*GE*_*ijk*_). This enables SINN to avoid the adverse effects of multimodal optimization of DNNs, while still utilizing all information in the prediction of genotype-environment interactions.

### Dataset

We used the Genomes to Fields 2022 Maize GxE prediction competition (G2F2022) data (Lima *et al*. 2023) and prediction setting in this work, to enable comparison between our approach with existing methods and competition results. The competition challenged participants to predict the plot yields of 505 new and 43 previously observed maize hybrids in 26 environments across North America for the year 2022, based on a training set of the plot yields of 4,683 maize hybrids in 217 environments across 47 locations in the years 2014-2021. The complete dataset included over 140,000 measured plots, from which 10,290 plots are assigned to the test set.

The dataset included genetic markers, weather and soil measurements, management information and plot yields. For each hybrid, information on 437,214 single nucleotide polymorphisms (SNPs) was provided. For each environment 29 soil features, 5 management features and 16 daily weather features were provided. Each plot record included planting date, harvest date and planting density. For a complete description of the dataset, we refer to Lima *et al*. (2023).

### Data preprocessing

The training dataset was filtered and imputed to ensure each training sample included a full set of features. Training samples without plot yields, genotypes without SNPs and trials without weather data were removed, leaving 123,517 samples for 4,417 genotypes and 212 environments, across 47 locations in 8 years. All features with a higher than 30% rate of missing values were removed. SNPs with a minor allele frequency below 1% or with more than 10% missing values were removed, and the resulting SNPs were randomly downsampled to 20,000 SNPs for the DNN-based methods. Imputation methods varied per feature, as follows: SNPs were imputed with the most frequent genotype at each locus. Management features consisted of two categorical features, representing whether the environment was irrigated and was a disease trial. For each, missing values were imputed with the most common class. Soil features were imputed by nearest spatial neighbor across training environments. Weather features within a growing season were imputed by linear interpolation in time. For each plot, weather features were aligned from 7 days before sowing to 133 days after sowing, giving a time series of 140 days for each plot. Missing values for planting date, harvest date and planting density were imputed with means within each environment where available, and global means elsewhere. The final preprocessed dataset retained 20,000 genetic markers, 11 daily weather features over 140 days, 20 soil features and 2 management features.

### Experiments

In order to investigate the proposed SINN framework, we conducted two experiments in the two main steps of the framework: modeling of components of yield, and structured prediction of yield. An overview of our experimental framework is shown in Figure 2. We split the dataset into two subsets: a cross-validation set (years 2014-2021) and a holdout test set (final year 2022). The cross-validation set was used for all statistical decompositions, model training and model cross-validation. The holdout test set was used only to evaluate trained models, and was identical to the test set used in the competition.

**Figure 2.**
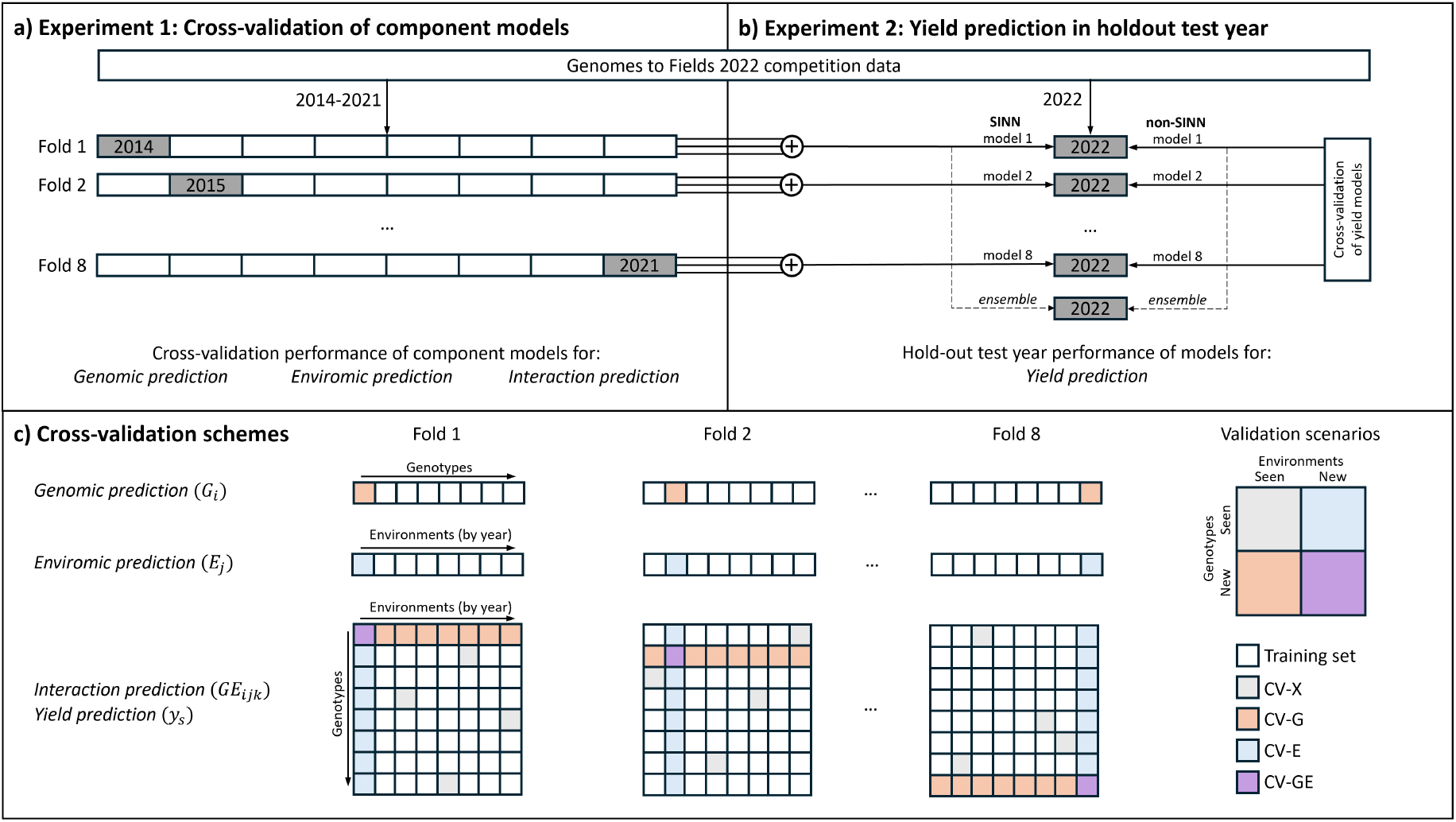
Overview of experiments and cross-validation schemes implemented in this work. a) Experiment 1. In the cross-validation set (years 2014-2021), we performed eight-fold cross-validation of component models for genomic prediction, enviromic prediction and interaction prediction. b) Experiment 2. We evaluated yield prediction models in the holdout test set (year 2022), measured stability in test set performance over cross-validation training sets (model 1-8), and measured performance of averaged predictions (ensemble). SINN models were constructed through summation of component models from Experiment 1, non-SINN models were fitted directly on yield. c) Cross validation schemes and scenarios for all modeling tasks defined in Experiment 1 and 2.

#### Experiment 1: Cross-validation of component models

In our method, we proposed to model unimodal (genetic, environmental) and multimodal (interaction) components of yield separately. This created the opportunity to evaluate modeling approaches per component task. We applied the decomposition in (3) to the cross-validation set (years 2014-2021) and obtained unimodal genetic targets *G*_*i*_, unimodal environmental targets *E*_*j*_ and residual genotype-environment interaction targets *GE*_*ijk*_. For each type of component task with corresponding target, we evaluated a set of models across eight cross-validation folds. The set of component models used per component target are further described in section *Models*.

As illustrated in Figure 2c, different cross-validation schemes were used for each component task. Genomic prediction of *G*_*i*_ was evaluated on new genotypes, corresponding to scenario CV-G. Each validation fold for CV-G contained 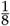 of genotypes. Enviromic prediction of *E*_*j*_ was evaluated on new environments, corresponding to scenario CV-E. We split environments based on a single year per validation set, again obtaining eight folds.

For interaction prediction of *GE*_*ijk*_, we defined a cross-validation scheme with four scenario. For each fold, 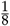 of genotypes and one year of environment were selected, corresponding to the same validation genotypes and environments used for *G*_*i*_ and *E*_*j*_. These settings formed cross-validation scenarios CV-G and CV-E. Their intersection formed scenario CV-GE, predicting interactions for new genotypes and environments. Finally, we obtained a fourth validation scenario by sampling random combinations of observed genotypes and observed environments, and holding out all replicates of these combinations in the validation fold. This reflects scenario CV-X, new interactions of observed genotypes and environments. We sampled 2000 interactions for each fold of CV-X. Model performance in scenario CV-GE was used as target criterion for model and hyperparameter selection, as it reflects most closely the scenario of the holdout test set.

#### Experiment 2: Yield prediction in holdout test year

In order to evaluate the effectiveness of the proposed SINN method, we conducted a second experiment that mirrored exactly the test conditions of the Genomes to Fields 2022 competition (Washburn *et al*. 2025). We evaluated a set of models on a holdout test set (year 2022), comprising of 43 observed and 505 new genotypes grown in 26 new environments. All environments were in repeated locations from previous years, but had new weather conditions.

Models were evaluated in their capacity to predict maize yields (Mg/ha) of genotype-environment combinations. The competition reported a single target yield value per combination, which we infer is an average over an unknown number of replicates. We constructed the complete SINN model from component SINN models evaluated in *Experiment 1*, following (6). This is shown in Figure 2b as summation of three component models per cross-validation fold.

Non-SINN models (further specified in *Models*) were not constructed from component models in *Experiment 1*. Instead, they were directly fitted on observed plot yields in the cross-validation set. In order to correctly compare variability in model errors between SINN and non-SINN methods, we fitted one model of each non-SINN model type per cross-validation fold, with the same cross-validation scheme used for interaction prediction. Again, cross-validation scenario CV-GE was used for model selection and hyperparameter optimization.

Using the models obtained through cross-validation would have underestimated the potential performance of models on the holdout test set, as each model was trained with a subset of all available samples. However, it is highly informative of stability of modeling approaches across dataset subsets. To retain this information on stability while avoiding under-estimation of model performance, we reported both predictions of individual models and ensembled predictions per model. We reported performance of individual models when comparing models which we implemented in this study, but reported ensembled predictions when comparing with current state-of-the-art from Washburn *et al*. (2025). For each model type, we ensembled predictions for the hold-out test set over the set of eight models obtained through cross-validation.

### Metrics

As main evaluation metrics we used the Root Mean Squared Error (RMSE), Mean Absolute Error (MAE) and the Pearson correlation coefficient (*r*). The Mean Squared Error (MSE) was used as optimization criterion. To compare prediction of genetic versus environmental signals, we define the *genetic accuracy* as average within-environment correlation, calculated as following:

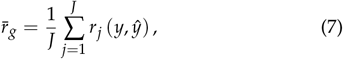

Similarly, we define the *environmental accuracy* as average within-genotype correlation:

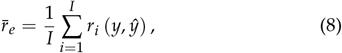

### Models

We used a similar approach for fitting each component of SINN, 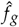,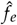 and 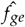. The models were fitted with the AdamW optimizer (Loshchilov and Hutter 2019), minimizing Mean Squared Error (MSE) as training objective. We linearly increased the learning rate from zero to the chosen maximum learning rate over the first 5 training epochs of each model. The number of training epochs for each task was determined by manual experimentation prior to hyperparameter tuning. The final validation fold was used for hyperparameter tuning of each model. For environments, this fold corresponded to the trials conducted in 2021. The search space and number of iterations per model of the hyperparameter selection are described in Appendix A.

As genomic prediction model *f*_*g*_ we used a multi-layer perceptron (MLP) (Rosenblatt 1958). Input SNPs were encoded as the count of alternate alleles minus 1 (−1, 0 or 1), resulting in a genetic input dimension *d*_*g*_ of 20,000. The MLP used had 3 layers: one input layer, one hidden layer and one output layer with 96, 48 and 1 nodes respectively. Each input and hidden layer was followed by a layer normalization (Ba *et al*. 2016) and dropout layer (Srivastava *et al*. 2014), then an activation function. We used a ReLu layer (Nair and Hinton 2010) as activation function. Before the output layer, a sigmoid activation function was used instead. We trained *f*_*g*_ for 250 epochs, with a batch size of 256, a learning rate of 0.01 and a weight decay of 1 × 10^−4^.

As enviromic prediction model *f*_*e*_ we used the same base architecture as *f*_*g*_. Daily weather features were averaged over the growing season and concatenated with soil and management features, resulting in an environmental input dimension *d*_*e*_ of 33. The model had one input layer, two hidden layers and one output layers with 64, 64, 32 and 1 nodes respectively. We trained *f*_*e*_ for 500 epochs, with a batch size of 32, a learning rate of 0.001 and a weight decay of 3 × 10^−5^. We noted that the initial performance of *f*_*e*_ during hyperparameter tuning was unstable and resulted in models with a low Pearson Correlation Coefficient, but low error. This indicated selection for models which capture the validation set mean yield, but not structure. To rectify this, we added a penalty of − 5*r* to the criterion, resulting in a pragmatic selection criterion of *MSE* − 5*r* with approximately equally scaled MSE and *r*.

As interaction prediction model, we reused the trained 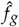 and 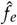 as encoder networks. The output of the final linear hidden layer of each encoder was projected with a learned linear projection to an embedding vector of size 64. To limit our model to first-order interactions, the genetic and environment embeddings were fused using a dot product operation to obtain predicted yield values. Training consisted of learning the linear projection weights and finetuning the parameters of the encoder networks. We trained the model for 250 epochs, with a batch size of 256, a learning rate of 0.01 and a weight decay of 3 × 10^−3^. In this work, we denote all models that are constructed using *f*_*g*_, *f*_*e*_ and *f*_*ge*_ with *SINN*, as they were constructed through the SINN framework. However, we note that models that exclude *f*_*ge*_ do not allow for interactions, making the *I* superfluous. Nonetheless, we define the following set of SINN models:

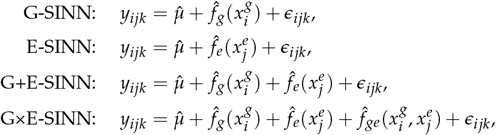

Throughout the results, G×E-SINN and SINN are used interchangeably. Additionally, SINN-based methods without *f*_*ge*_ (G-SINN, E-SINN, and G+E-SINN) should be understood as ablations of the SINN framework, not as separate modeling approaches.

### Baseline models

As baseline models, we used the winning model and the highest ranking DNN of the G2F2022 dataset competition (Washburn *et al*. 2025). Additionally, we fitted a set of commonly utilized statistical reaction-norm-based models that extend G-BLUP (Meuwissen *et al*. 2001) with an environmental covariance matrix and interaction matrix based on genetic markers and environmental covariates (Jarquín *et al*. 2014). In contrast to the models fitted in the original work, we did not include separate (unstructured) effects for lines and trials, to avoid absorption of effects from marker and environmental covariate-based terms. We define this set of models as following: with components defined as following:

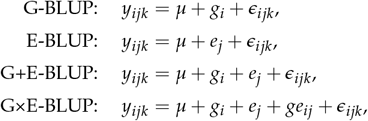

with components defined as following:

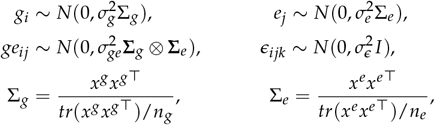

where again *y*_*ijk*_ denotes yield for genotype *i*, environment *j* and replicate *k, g*_*i*_ is the predicted genomic effect, *e*_*j*_ the predicted effect of the environment, and *ge*_*ij*_ the predicted effect of the genotype-environment interaction. Σ_*g*_ was the genomic relationship matrix of *n*_*g*_ genotypes, Σ_*e*_ the environmental relationship matrix of *n*_*e*_ environments, calculated from genetic features *x*^*g*^ and environmental features *x*^*e*^ respectively. For *Experiment 2*, we fitted each BLUP-based model on yield directly. Memory requirements for Σ_*g*_ ⊗ **Σ**_*e*_ scaled quadratically with number of samples. As our experiments were constrained to 500GB of RAM, we were forced to drop replicates and entries for the highest frequency genotypes until 40% of the samples remained when fitting G×E-BLUP. To keep implementation consistent between BLUP-based models, we used the same downsampling fraction as G×E-BLUP for other BLUP-based models fitted on yield. Initial experiments indicated that predictive performance of all BLUP-based methods saturated at downsampling fractions below 40%. As such, we would expect no increase in performance if the downsampling fraction were increased beyond 40%. For *Experiment 1*, we fitted one BLUP-based model on each set of labels: we fitted G-BLUP on *G*_*i*_, E-BLUP on *E*_*j*_, and G×E-BLUP on *GE*_*ijk*_. Again, we dropped replicates from *GE*_*ijk*_ until 40% of samples remained.

As additional DNN baseline, we replicated a recent DNN-based model by Kick *et al*. (2023) that was developed using a subset of the G2F2022 dataset. We only tuned the learning rate and weight decay. The model architecture and other hyperparameters remained unchanged. The model consisted of a genetic encoder, a soil encoder, a weather encoder and an interaction module. The genetic encoder was an MLP with one hidden layer using principal components of *x*^*g*^ as inputs. The weather encoder was a Convolution Neural Network (CNN) taking daily weather values as inputs. The soil encoder was an MLP with one hidden layer taking the soil and management values as input. The interaction module took the concatenated outputs of the final hidden layer of each encoder as input and was an MLP with five hidden layers. For further details, we refer to the original implementation (Kick *et al*. 2023).

For the first set of experiments, we trained only the relevant DNN components, adding a linear output layer if not present. We trained the genetic encoder on *G*_*i*_ for 250 epochs, with a batch size of 256, a learning rate of 3 × 10^−4^ and a weight decay of 1 × 10^−4^. We jointly trained both the weather and soil encoders on *E*_*j*_ for 500 epochs, with a batch size of 32, a learning rate of 1 × 10^−3^ and a weight decay of 3 × 10^−4^. To predict *GE*_*ijk*_, we followed the same procedure as the SINN model, and initialized the full DNN with the trained encoder weights. Then, we trained the model on *GE*_*ijk*_ for 250 epochs, with a batch size of 192, a learning rate of 3 × 10^−3^ and a weight decay of 3 × 10^−3^. For the second set of experiments, we trained the full model with yield as target variable on CV-GE. Following hyperparameter tuning, we trained for 150 epochs with a batch size of 192, a learning rate of 0.01 and a weight decay of 3 *×* 10^−3^.

## Results

In this section, we provide a stagewise analysis of the proposed SINN framework. Following the modeling steps of SINN (decomposition, component modeling, structured prediction), we focus on answering the following three questions:

### Preliminary

How is the variance in yield partitioned among genotypes, environments and genotype-environment interactions?

### Experiment 1

How do different types of models predict isolated unimodal and multimodal components of yield?

### Experiment 2

How well does SINN predict maize yield in the next year of a multi-environment trial, in comparison to current state-of-the-art?

### Preliminary: Analysis of variance

As SINN relies on a statistical decomposition of yield across genotypes, environments and genotype-environment interactions, we preclude our results with an analysis of variance for the data used in model fitting and cross-validation (years 2014-2021). The current implementation of SINN utilizes an additive decomposition. We use a different, more informative decomposition to investigate the variance of yield across the training data. We fit a three-way random effect ANOVA between genotype, year and location, which is shown in Table 1.

**Table 1.**
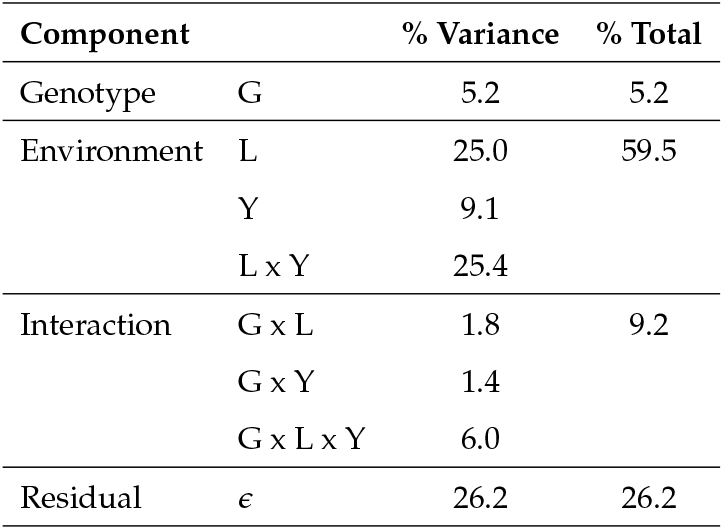
Three-way ANOVA of yield by genotype (G), location (L) and year (Y) for cross-validation set spanning 2014-2021.

Genotype, environment and interaction components contributed 5.2%, 59.5% and 9.2% to overall variance. The location effect contributed 42% of the environmental variance, suggesting a considerable location-specific component to environmental effects. Finally, the residual variance was 2.85 × the genotype-by-environment variance, indicating that the genotype-by-environment specific signal is challenging to separate from the residual within-environment noise.

### Experiment 1: Cross-validation of model components

We now consider the performance of component models in predicting the decomposed yield components *Ĝ*_*i*_, *Ê*_*j*_ and *ĜE*_*ijk*_ for cross-validation folds across the 𝒮. An overview of the results of each component is shown in Figure 3, with colors matching model types (BLUP-based, vanilla DNN, and SINN). For reference, cross-validation schemes per component task were provided in Figure 2c.

**Figure 3.**
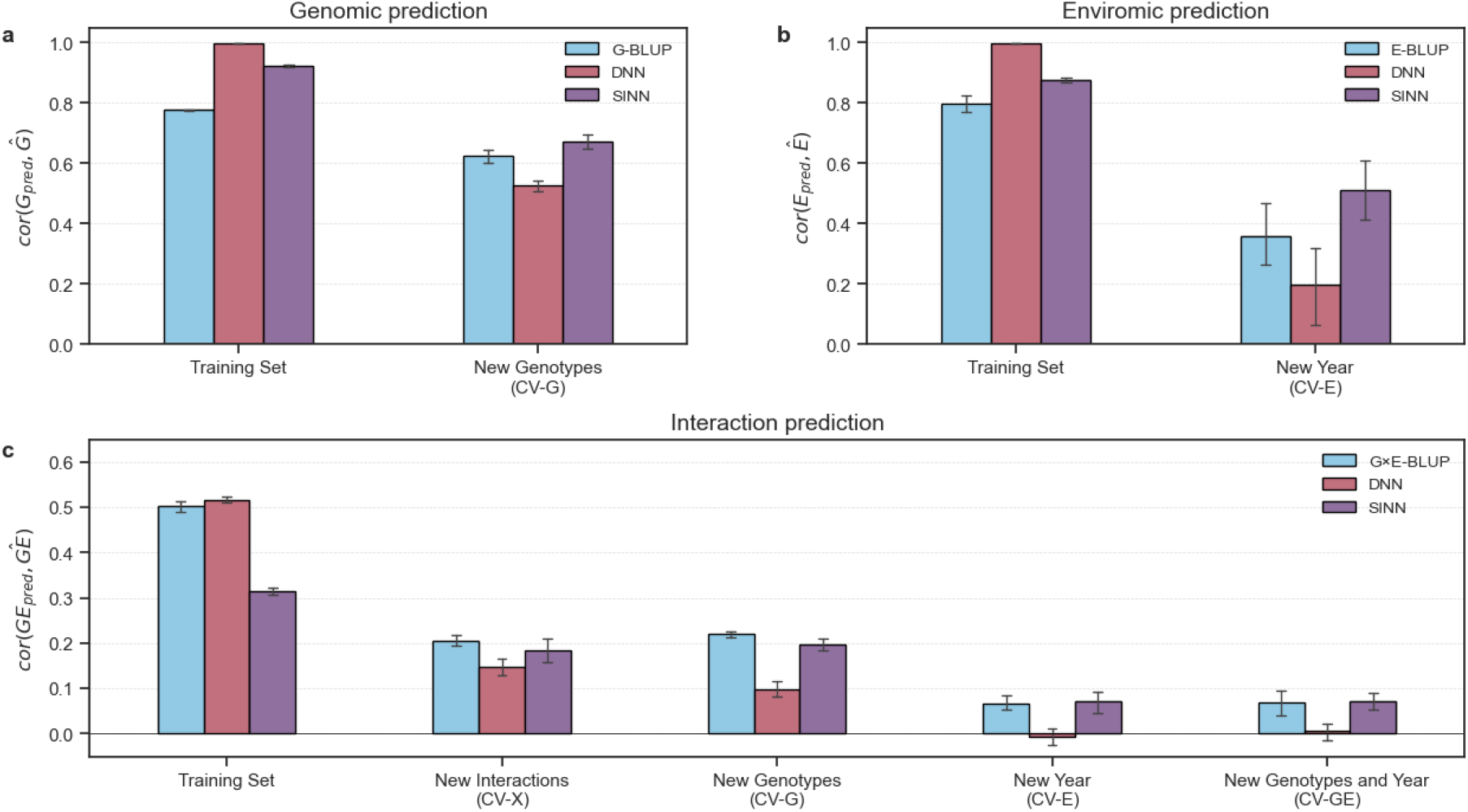
Predictive performance of SINN component models (purple) compared to BLUP-based (blue) and DNN (red) component models. Error bars show standard deviation over 8 cross-validation folds. (a) Predictive ability for genomic prediction of the estimated effect of new genotypes (CV-G). (b) Predictive ability for enviromic prediction of new environments (CV-E). (c) Predictive ability for interaction prediction of residual genotype-by-environment interaction in scenarios from left to right: training set, new interactions of observed genotypes and environments (CV-X), new genotypes and observed environments (CV-G), observed genotypes and new environments (CV-E), and new genotypes and environments (CV-GE).

First, the genomic prediction results for genetic effects *Ĝ*_*i*_ of new genotypes (CV-G) are shown in Figure 3a. The genetic encoder of SINN (MLP) had the highest cross-validation predictive ability, followed by G-BLUP and the genetic encoder of DNN (principal component-based MLP). Furthermore, the baseline DNN had a correlation of near 1 on the training set, indicating severe overfitting.

We show the enviromic prediction results of environment effects *Ê*_*j*_ in new environments (CV-E) in Figure 3b. Overall, the cross-validation predictive ability was lower than for genomic prediction and showed a larger variance over folds. The enviromic encoder of SINN (MLP) had the highest performance, followed by E-BLUP and the environment encoder of the DNN (CNN). The baseline DNN had a near-perfect training correlation, again indicating severe overfitting.

Finally, the prediction of residual genotype-by-environment interactions *ĜE*_*ijk*_ is shown across four validation scenarios in Figure 3c. The performance of all models was highest when predicting new interactions for observed hybrids and observed trials (CV-X). Surprisingly, G×E-BLUP and SINN showed no decrease in predictive ability when predicting for new genotypes (CV-G) compared to CV-X, indicating strong generalization of performance to new genotypes. The DNN dropped in performance from an predictive ability of 0.15 to 0.10 for new genotypes, indicating overfitting of interactions on observed genotypes.

All three models showed a large drop in performance when predicting for new environments (CV-E) compared to CV-X, indicating poor generalization of interaction prediction to new environments. On the cross-validation scenario reflecting the G2F2022 test scenario, predicting interactions for new genotypes and environments (CV-GE), SINN and G×E-BLUP had a similar predictive ability of 0.07, while the baseline DNN had a predictive ability of 0.01.

For all models, the predictive ability of CV-GE was similar to CV-E, but considerably lower than CV-G, indicating that the low reported accuracies for CV-GE are primarily driven by poor generalization to new environments. Furthermore, the three-way ANOVA decomposition in Table 1 can be used to estimate the predictive ability of a perfect interaction prediction model. The interaction-specific variance divided by the total interaction plus residual variance is estimated as 9.2/35.4 ≈ 0.26. As such, we can roughly attribute 26% of the variance in *ĜE*_*ijk*_ to genotype-environment interactions. Converting from explained variance to correlation, we obtain a maximum achievable correlation of 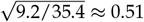 for a model that perfectly predicts the genotype-environment interactions for each genotype-environment combination. Both G×E-BLUP and DNN reached this upper bound on the training set, which again indicates overfitting. Furthermore, this irreducible error lowered the observed predictive ability of all interaction prediction models across all scenarios, as it includes the accuracy of predicting residual noise.

Overall, SINN exhibited lower overfitting and better generalization across all tasks and scenarios when compared to the baseline DNN components. SINN showed the largest improvements over BLUP-based models in predicting *Ê*_*j*_, a smaller improvement in predicting *Ĝ*_*i*_ and a comparable performance in predicting *ĜE*_*ijk*_. Interaction prediction was the most challenging task out of the three component tasks, followed by enviromic prediction and genomic prediction. Difficulty of interaction prediction scenario followed the same trend, with lower predictive abilities reported for prediction in new environments (CV-E) than for new genotypes (CV-G) across all models.

### Experiment 2: Maize yield prediction in holdout test year of a multi-environment trial

In the previous section, we observed that SINN outperforms other models on component modeling tasks. Next, we investigate whether this leads to an increased predictive ability for the prediction of maize yield in a holdout test year of a multi-environment trial. To this end, we compare SINN with other models fitted directly on yield, and with previously reported models in the The Genomes to Fields 2022 Maize Genotype by Environment Prediction Competition Washburn *et al*. (2025).

In Figure 4a the performance of three classes of models is shown in predicting maize yield in the holdout test year (2022) of the G2F maize hybrid dataset. We compare four statistical models including G-BLUP (Meuwissen *et al*. 2001) and three extensions including environmental covariates (E-BLUP, G+E-BLUP) and genotype-by-environment interactions (G×E-BLUP) following Jarquín *et al*. (2014). As baseline Deep Neural Network (DNN), we use the network architecture from Kick *et al*. (2023). We report on four variants of the SINN model: a genetic model component (G-SINN), an environment model component (E-SINN), an additive combination of the genetic and environmental model components (G+E-SINN), and a full model with genetic, environment and genotype-by-environment interaction (SINN). SINN had a higher and more consistent predictive ability (0.59±0.03) than DNN (0.38±0.07) and G×E-BLUP (0.39±0.07), which had approximately equal predictive ability. G-BLUP (0.15), E-BLUP (0.28) and G+E-BLUP (0.34) had a lower predictive ability than G-SINN (0.24), E-SINN (0.53) and G+E-SINN (0.58) respectively.

**Figure 4.**
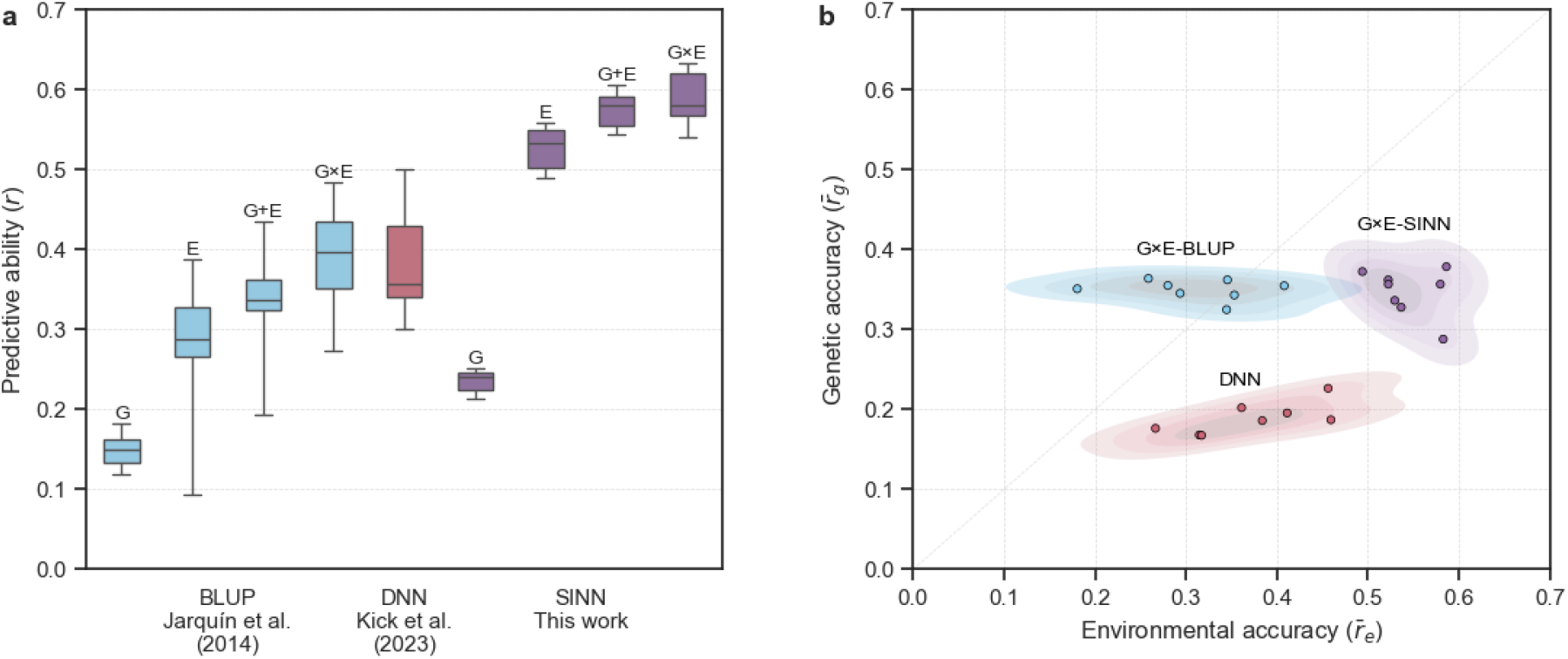
Predictive performance for plot-level maize yield in test year (2022) of Genome to Fields, including unobserved hybrids. (a) Overall predictive ability (Pearson Correlation Coefficient) of models fitted in 8 cross-validation folds. (b) Kernel density estimate of correlation per genotype and per environment of the same 8 models for G×E-BLUP (blue), DNN (red) and SINN (purple).

Dissecting overall predictive ability into genetic and environmental accuracies reveals qualitative differences in the predictions of BLUP-based, DNN-based and SINN-based models. Genetic and environmental accuracies, defined in (7) and (8), for the three models types with genotype-by-environment interactions are shown in Figure 4b. While G×E-BLUP and DNN had similar overall predictive ability, DNN had a significantly lower genetic accuracy (0.19) than G×E-BLUP (0.35), indicating it performed worse at predicting the relative performance of genotypes within each environment. DNN had a higher environmental accuracy (0.37) compared to G×E-BLUP (0.31). SINN matched G×E-BLUP in genetic accuracy (0.35), while improving over both G×E-BLUP and DNN in terms of environmental accuracy (0.54). Overall, SINN improves over DNN in terms of genetic and environmental accuracies, and improves over G×E-BLUP in terms of environmental accuracy.

Table 2 directly compares the three models with the competition results from Washburn *et al*. (2025). To keep models and data splits consistent across experiments, we did not fit a model using the full training and validation. Instead, we ensembled the predictions of models fitted on each cross-validation fold by direct averaging to obtain model predictions that incorporate information of the full training and validation set. Table1, reports standard deviation of results of individual models across scenarios within the test set: New Year (CV-E), and New Genotypes and Year (CV-GE).

**Table 2.**
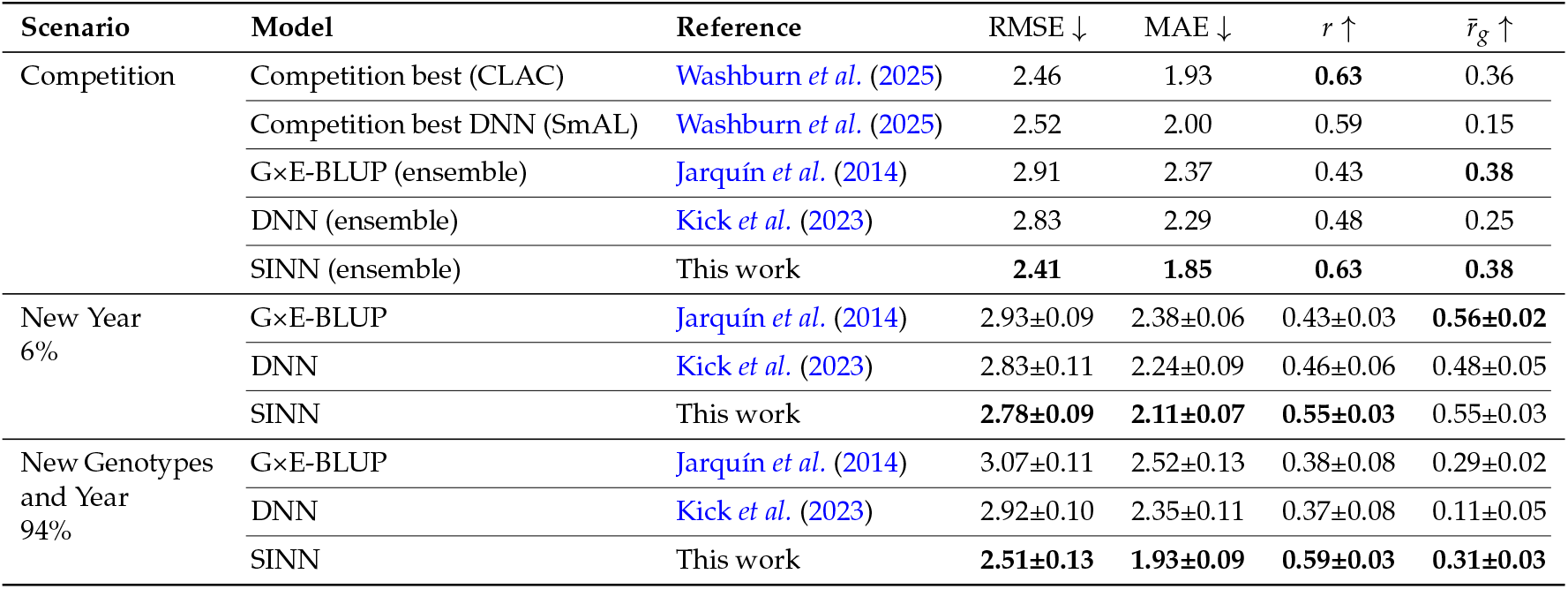
Predictive performance across test scenarios. Competition comprises the full test set, composed of 6% of samples with new environments in a new year, and 94% with both new genotypes and new environments in a new year. Best model per scenario is shown in bold. Standard error of models is shown as ±. For comparison with the G2F2022 competition, models are ensembled by averaging predictions.

**Table 3.** Settings and search spaces for hyperparameter selection

| Model | Target | # | Layers | Nodes per Layer | Learning Rate | Weight Decay |
| --- | --- | --- | --- | --- | --- | --- |
| G-SINN | $\hat{G}_i$ | 125 | | {64, 96, 128, 192, 256} | {1e-4, 3e-4, 1e-3, 3e-3, 1e-2} | {1e-4, 3e-4, 1e-3, 3e-3, 1e-2} |
| E-SINN | $\hat{E}_j$ | 200×5 | { 3, 4, 5 } | {8, 16, 32, 48, 64} | {1e-5, 3e-5, 1e-4, 3e-4, 1e-3} | {1e-5, 5e-5, 1e-4, 3e-4, 1e-3} |
| G×E-SINN | $\hat{G}_{E_{ijk}}$ | 125 | | {8, 16, 32, 64, 128 } | {1e-4, 3e-4, 1e-3, 3e-3, 1e-2} | {1e-4, 3e-4, 1e-3, 3e-3, 1e-2} |
| DNN (G encoder) | $\hat{G}_i$ | 25 | | | {1e-4, 3e-4, 1e-3, 3e-3, 1e-2} | {1e-4, 3e-4, 1e-3, 3e-3, 1e-2} |
| DNN (E encoder) | $\hat{E}_j$ | 25×5 | | | {1e-5, 3e-5, 1e-4, 3e-4, 1e-3} | {1e-5, 5e-5, 1e-4, 3e-4, 1e-3} |
| DNN | $\hat{G}_{E_{ijk}}$ | 25 | | | {1e-4, 3e-4, 1e-3, 3e-3, 1e-2} | {1e-4, 3e-4, 1e-3, 3e-3, 1e-2} |
| DNN | $y_s$ | 25 | | | {1e-4, 3e-4, 1e-3, 3e-3, 1e-2} | {1e-4, 3e-4, 1e-3, 3e-3, 1e-2} |

SINN obtained the lowest RMSE and MAE of all models. G×E-BLUP and SINN jointly had the genetic accuracy 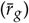. Similar to DNN, the competition best DNN had very low genetic accuracy, indicating that state-of-the-art DNNs for genomic prediction in new environments suffer do not predict genetic differences as well as BLUP- or SINN-based methods. SINN outperformed G×E-BLUP and DNN across both New Year (CV-E) and New Genotypes and Year (CV-GE) scenarios for RMSE, MAE and *r*. G×E-BLUP obtained a higher genetic accuracy in CV-E, while SINN obtained a higher genetic accuracy in CV-GE.

## Discussion

### Structured optimization guides neural networks towards improved utilization of genetics

Both in our own baseline DNN as in the best DNN from the G2F2022 competition, we observed a low genetic accuracy of predicted yield values. This indicates a tendency of DNNs to focus on environmental information, and under-utilize genetic information in predicting yield. This is a recognized failure mode of multimodal deep learning models (**?**), where the model greedily focuses on a single modality of data. By separating the prediction of each unimodal signal, SINN avoids this problem and improves the utilization of genetic information. Surprisingly, SINN also improved the prediction of the unimodal effect of the dominant environmental modality, as shown by the improved genetic accuracy with respect to the baseline DNN. We attribute this to reduced overfitting of SINN compared to DNN. Overall, this enabled SINN to outperform all DNN-based baselines, even with a relatively simple model architecture.

The statistical decomposition of yield into yield components enables the design of targeted neural network components that match model complexity and regularization to the information content of each yield component. Without this targeted design the neural network components tended to overfit on both genetic and environmental information, as was shown by the high training accuracies and low validation accuracies of the baseline DNN components. With sufficient regularization, the nonlinear neural network components demonstrated an advantage over linear BLUP-based models for predicting holdout genetic and environmental effects. Surprisingly, this improvement was smallest for interaction prediction, which has both the highest number of samples as well as the expectation of being highly nonlinear. Across our interaction prediction experiments, we noticed that neural networks had a strong tendency to overfit and were limited to a very simple interaction component to succesfully generalize to new environments. We theorize that adding model complexity to any component downstream of the environmental information can push the model towards overfitting on the training environments, and that this occurs at lower interaction component complexity than overfitting on the training genotypes.

However, it is important to note that the decomposition of yield into components has two effects on the targets: it reduces noise in the components, and also introduces systematic errors from the statistical model assumptions which propagate to downstream functions. The first was observed by comparing the relative performance of BLUP-based models and baseline DNN-based models on predicting the differences between environments. On a per-component basis, E-BLUP outperforms the DNN environment encoder in predicting *Ê*_*j*_, while DNN outperforms G×E-BLUP in predicting differences between environments when trained directly on yield. Noise has a regularizing effect on DNN optimization, so DNN model components have a stronger tendency to overfit when this noise is removed. While the regularization of the DNN model was tuned through weight decay and learning rate, this was insufficient to avoid overfitting without modifying the network architecture. However, with full tuning of regularization and model complexity, this separation becomes beneficial, as it enabled SINNs to utilize genetic information more effectively than end-to-end optimized DNNs.

### Understanding the challenges of deep learning-based interaction prediction

One commonly stated aim for applying deep learning based methods to genomic prediction in new environments is to predict nonlinear genotype-by-environment interactions. However, this aim cannot be validated based current evaluation protocols, which rely on aggregate accuracy metrics evaluated across cross-validation scenarios.

SINN enables us to measure interaction prediction performance in isolation, and thus allowed us to directly test this aim for the first time. Surprisingly, we found that the baseline DNN from Kick *et al*. (2023) failed to predict any genotype-by-environment interactions for holdout environments, when interactions were isolated from the genetic and environmental main effects. When evaluated directly on yield, the additive structure of BLUP-based and SINN-based methods enable quantification of the contribution of predicted interactions. However, current evaluation protocols do not allow us to evaluate whether the DNN predicts any interactions in this context. For method development, a failure to measure progress means a failure to improve. SINN allows us to measure interaction prediction performance in isolation, and thus enables the targeted improvement of DNNs for interaction prediction within the framework. However, it does not enable the measurement of interaction prediction performance for models fitted end-to-end on yield. New metrics are needed for this purpose. As a potential diagnostic tool we identify the empirical multimodally-additive function projection (EMAP) by Hessel and Lee (2020), which measures black box function performance without predicted interactions.

### Prediction of environment-specific ranking is limited by number of independent environments

In our results, we compared interaction prediction performance of models across multiple scenarios and found that all models show a large decrease in performance when predicting in new environments. We also note that a low-complexity neural network interaction component was needed to limit overfitting on environments, even though the models were trained on over 100,000 interactions. This mirrored the results of predicting *Ĝ*_*i*_ and *Ê*_*j*_, as prediction performance for *Ê*_*j*_ was considerably lower than for *Ĝ*_*j*_, and a low-complexity neural network component was needed to avoid overfitting on *Ê*_*j*_. This comes as no surprise for *Ê*_*j*_, as only 217 unique environments were available in the training set, and DNNs typically need 1,000s to 10,000s of independent samples to improve over other methods.

Surprisingly, a similar overfitting pattern was observed for the 100,000 interaction prediction samples. A potential explanation is that the interactions are not independent samples, but repeated realizations of the same 217 environments, which encourages overfitting on environmental information. This suggests that the main limitation to prediction of environment-specific genetic rankings in new environments is the limited number of unique environments, not the number of unique genotypes. This is consistent with findings from Washburn *et al*. (2021), which found that yield prediction performance of DNNs could be improved by integrating a large number of unique environments, even when these environments had low-quality yield labels. Additionally, this suggests that DNN-based approaches for predicting genetic rankings in new environments might favor recently proposed sparse MET designs (Jarquin *et al*. 2020; Crespo-Herrera *et al*. 2021), where fewer genotypes are grown in each environment in favor of more unique realizations of environments.

### SINN can utilize advanced statistical models for multienvironment trials

Error propagation from the statistical decomposition is an unwanted by-effect of SINN. Any errors in the decomposition of *y*_*s*_ into *Ĝ*_*i*_, *Ê*_*j*_ and *ĜE*_*ijk*_ propagate through the training of *f*_*g*_, *f*_*e*_ and *f*_*ge*_ to downstream predictive ability. Addressing this error propagation completely would require integration between the statistical decomposition and the end-to-end optimization of DNNs, which we leave for further work. As a more practical approach, we propose to improve the decomposition of yield into *Ĝ*_*i*_, *Ê*_*j*_ and *ĜE*_*ijk*_ with statistical models for estimating genotype-environment interactions. In this work, we deliberately pursue a minimal implementation of SINN and thus utilize simple DNN components, and a minimally complex decomposition. However, as we note in section *Introducing SINN*, this naive decomposition obtains biased estimates of 𝔼 [*y* | *i*] and 𝔼 [*y* | *j*]. Fitting *G*_*i*_ and *E*_*j*_ as random effects could improve these estimates, as the MET in the G2F2022 dataset is imbalanced and incomplete. Furthermore, the current decomposition does not separate genotype-interaction from noise. This can be realized by a wide range of modern statistical models for both genotype-environment interaction and noise, including modeling of experimental design (blocks, rows, columns), spatial trends within trials with P-splines (Velazco *et al*. 2017; Rodríguez-Álvarez *et al*. 2017), and variance-covariance structures of genotype-environment interactions with modern factor-analytic models (Piepho and Williams 2024).

## Conclusion

We demonstrate the Structured Interaction Neural Network (SINN), a new framework for the optimization of deep neural networks (DNNs) for genomic-enviromic prediction. We show that SINN improves genomic prediction of maize yield in future environments when compared to a kernel-based statistical model, a specialized DNN, and the best models of an open benchmark dataset.

SINN reduces overfitting and improves prediction of genetic effects, environmental effects and genotype-by-environment interactions when compared to DNN models, and improves prediction of environmental effects when compared to kernel-based statistical models. Our findings further indicate that prediction of environment-specific rankings is mainly limited by generalization to new environments, rather than generalization to new genotypes. Surprisingly, we find that the baseline DNN had the lowest performance and failed to predict any genotype-environment interactions in new environments, when interactions are predicted in isolation.

Our SINN framework structures further MMDL research, as it decomposes both the task and model into components that can be improved independently. Additionally, it opens the door to further integration of advanced statistical decomposition methods with new DNN approaches. By bridging statistical genetics and deep learning, SINN provides a new approach to accurate, modular and scalable prediction of genotype-environment interactions.

## Data availability

All data used in this study is freely available from The Genomes to Fields 2022 Maize Genotype by Environment Prediction Competition, available on CyVerse under https://pubmed.ncbi.nlm.nih.gov/37461058/. Upon publication, the code and scripts used in this study will be made publicly available in the following GitHub repository: https://github.com/WUR-AI/sinn.

## Acknowledgments

The authors acknowledge the Genomes To Fields Initiative for providing the data utilized in this study.

## Funding

This work was partially supported by the Horizon Europe project PHENET - Tools and methods for extended plant PHE-Notyping and EnviroTyping services of European Research Infrastructures (Grant agreement ID 101094587).

## Conflicts of interest

The authors declare no conflicts of interest.

## Appendix Appendix A. Hyperparameter selection

